# Establishing BARA: Biological Nitrogen Fixation for Future Agriculture

**DOI:** 10.1101/684324

**Authors:** Muhammad Salar Khan, Naoru Koizumi, James L. Olds

## Abstract

The extensive use of nitrogen (N) fertilizers implicates a paradox. While fertilizers ensure the supply of a large amount of food, they cause negative environmental externalities including reduced biodiversity, eutrophic streams, and lakes. Moreover, such fertilizers may also result in a major public health hazard: increased antibiotic resistance. This *Perspective* discusses a critical role of perturbations in N cycle caused by excessive use of fertilizers and resulting implications as they relate to resistance genes and biodiversity in the biosphere. While there are solutions such as cover crops, these solutions are expensive and inconvenient for farmers. We advocate the use of biological fixation for staple crops—microbiome mediated natural supply of fixed N. This would involve engineering a microbiome that can be grown cheaply and at scale (less expensive than Haber-Bosch fertilizers). We also propose a practical framework of where and how research investments should be directed to make such a solution practical. We make three recommendations for decision makers to facilitate a successful trajectory for this solution. First, that future agricultural science seek to understand how biological fixation might be employed as a practical and efficient strategy. This effort would require that industries and government partner to establish a pre-competitive research laboratory equipped with the latest state-of-the-art technologies that conduct metagenomic experiments to reveal signature microbiomes. Second, the Department of Agriculture and state governments provide research and development (R & D) tax credits to biotech companies specifically geared towards R&D investments aimed at increasing the viability of biological fixation and microbiome engineering. Third, governments and the UN Food and Agriculture Organization (FAO) coordinate Biological Advanced Research in Agriculture (BARA)—a global agricultural innovation initiative for investments and research in biological fixation and ethical, legal, and social implications of such innovation.

## 1. Introduction

Earth’s human population is increasing. The current world population of 7.6 billion is projected to reach 9.8 billion in 2050 (UN *World Population Prospects*, 2017). For the ever-growing population, demand for food is rising. According to the Food and Agriculture Organization of United Nations (FAO), Food production must increase by 70% to feed a larger population (FAO *How to Feed the World 2050*, 2009). At this moment, the planet remains far from attaining the *Sustainable Development Goal 2* of *Zero Hunger*, which pledges to *end hunger, achieve food security, improve nutrition, and promote sustainable agriculture*. About 821 million people, i.e., one in nine, suffer from malnutrition across the world (WFP, 2019). This situation is more prevalent in developing countries. One in four are undernourished in Sub-Saharan Africa alone (Food Aid Foundation, 2018). The Earth’s human population is 2B over its natural carrying capacity (Erisman et al. 2008)—a result of the “Green Revolution” and the ability to chemically synthesize N-based fertilizers.

The practical application of chemical fertilizers to agricultural food systems is far from 100% efficient. In this *Perspective*, we focus on the policy issues related to surplus fixed N which pollutes freshwater bodies as well as estuarine waters, thereby contributing to eutrophication and significantly reducing biomass and biodiversity (Lewis et al., 2011). Results of a recent analysis suggest that the number of oceanic “dead zones” caused by eutrophication will increase dramatically (Altieri and Geden, 2014). Other than negative environmental impact, excessive N fertilizers have been linked to both the prevalence of antimicrobial resistance genes (ARGs) in animals and antibiotic-resistant bacteria (ARB) in humans (Liu et al., 2017; Singer et al., 2016). As microbes increase their resistance to the current antibiotic medicines, humans are increasingly at risk of developing infections for which there is no effective treatment.

### The extensive use of N fertilizers points to a paradox

while fertilizers ensure the supply of a large amount of food, they cause negative environmental externalities including reduced ecosystems services and a major public health hazard: the increased prevalence of antibiotic resistance in clinically relevant microbial species.

This policy *Perspectiv*e focuses on the critical role of perturbations of the N cycle caused by excessive use of fertilizers, resulting in negative outcomes both to the biosphere and human society embedded within it.

To reduce some of the damage caused by fertilizers’ overuse, Farina and colleagues (2018) advocated for improved crop rotation. Similarly, Dong et al. (2012) propose moderating application of fertilizers following a robust soil testing that accurately predicts N needs. While these proposed policy solutions are promising, they are expensive and inconvenient for farmers.

Here, we present biological fixation (microbiome mediated natural supply of fixed N to plants) for the major commercially relevant staple crops (maize, rice and wheat). Our view is that the microbiome mediated biological supply of N to plants is a readily available, cost-effective solution that is both replicable and applicable on a vast scale. We advocate the use of metagenomics and synthetic biology to essentially engineer a microbiome that can be manufactured cheaply and at scale (less expensive than Haber-Bosch fertilizers). To this end, we propose a practical framework of where and how the research should be headed to establish the viability and validity of this solution. Finally, we recommend a set of policies to implement a coherent government-wide investment strategy to ensure that future agriculture is sustainable, cheap, and sufficient.

## 2. Nitrogen use and overuse in agriculture lead to a nitrogen gradient

According to a UN FAO report (2004), Haber-Bosh derived N input dominates fertilizer supplementation for most cereal crops in the ecosystem. The report attributes the success of the “Green Revolution” to the significant role played by N fertilizers. A recent study published in PLoS Biology suggests that an overall production resulting from the use of N fertilizer represents 1-2% of global energy use (Van Deynze et al., 2018).

With increasing agricultural activity, gross annual N fertilizer use will likely increase (UN FAO, 2004). Lamarque and colleagues (2005) assert that anthropogenic N inputs are expected to more than double globally over the next century. Recent evidence provided by Li et al. (2016) already points to a substantial N increase in sections of the Midwestern United States, resulting from increased agricultural activity (Figure 1). Based on N deposition data readily available at the National Atmospheric Deposition Program (NADP), we displayed on a map variation of estimated N across the NADP sites. What we see is a striking continental-scale N gradient that correlates with agricultural crop production in the central United States (Baker et al., 2011). Agricultural crop activity map in figure 2 correlates with figure 1 as shown below.

**Figure 1:**
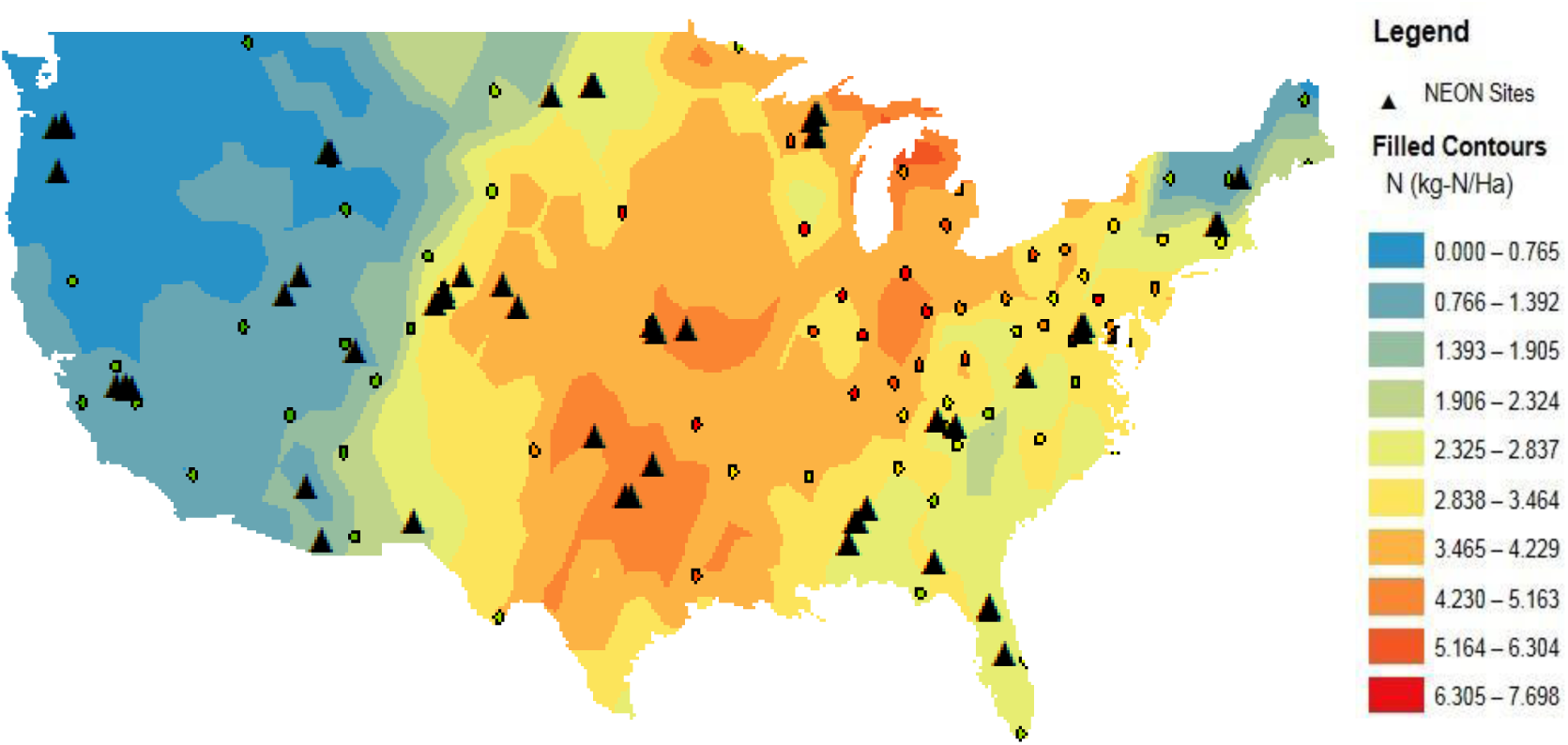
CONUS distribution of N for 2016 based on observations from the National Atmospheric Deposition Program. National Atmospheric Deposition Program sites are shown as color-coded points. The black triangles indicate National Ecological Observatory Network sites, which are strategically located sites across the United States to capture the variance in ecological and climatological measurements. The color at each point reflects the N value at the National Atmospheric Deposition Program site. The deeper red colors correspond to high values of N, whereas blue represents lower values of N. The background of the map displays continental variation in estimated N based on N values available for sites of the National Atmospheric Deposition Program.

**Figure 2:**
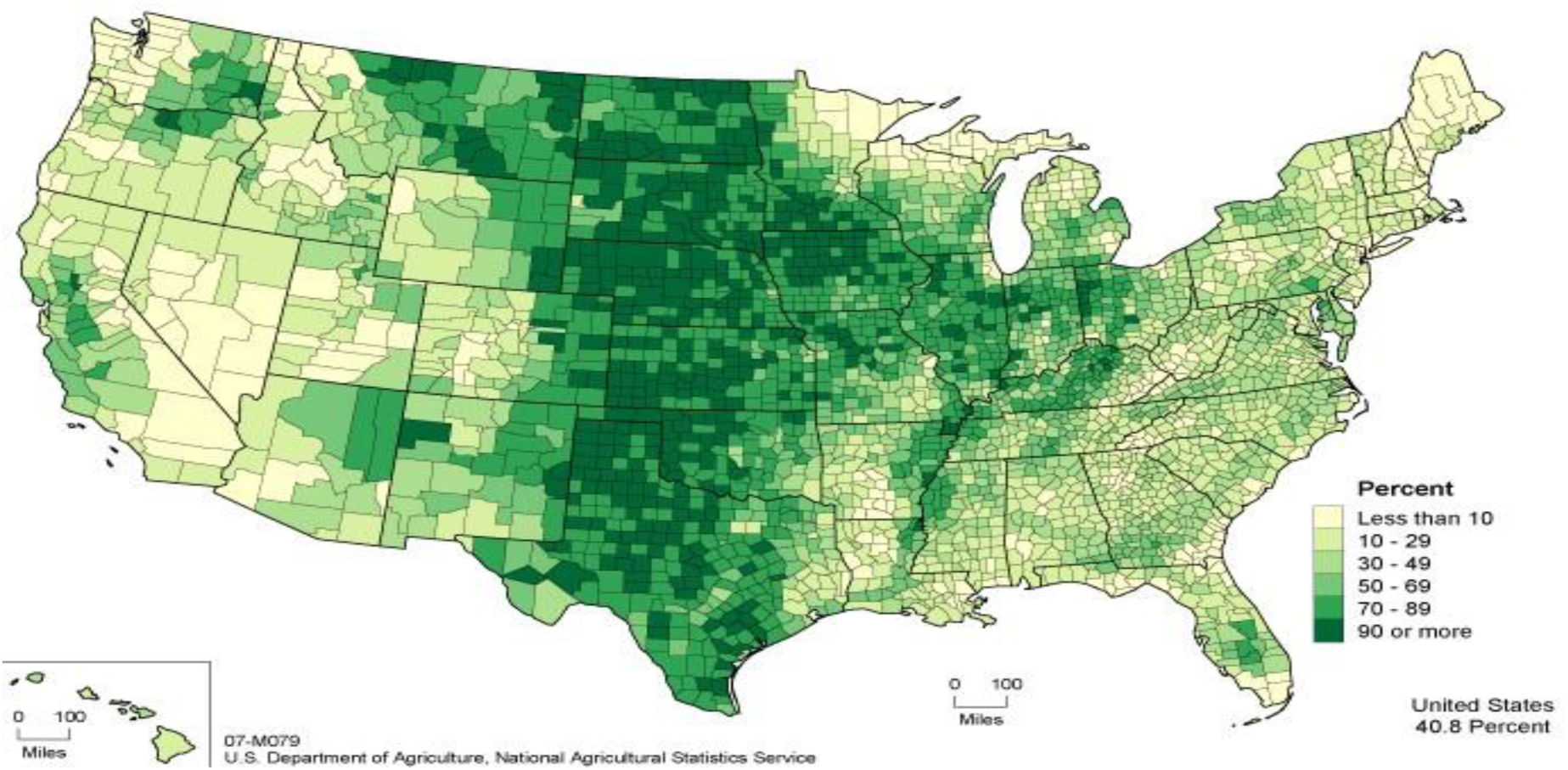
Acres of land in farms as percentage of land areas in acres (2007). The dark green areas are agriculture-intensive areas. While the light green and yellow areas are relatively low agriculture-intensive areas. The map shows continental variation in agricultural activity across the US. Figure credit: USDA. Figure taken from: https://scied.ucar.edu/sites/default/files/images/large_image_for_image_content/acres_land_farms.gif

## 3. Importance of nitrogen for life

As one of the building blocks of life, N is essential for the survival of plants, animals, and humans. It is a universally occurring, central chemical constituent of the biosphere, primarily derived from its fundamental role in nucleic acid and protein biochemistry. Plant tissues contain about 1.5% N in their total dry weight (Epstein, 1999). Since N is the constituent of many plant cell components (DNA, RNA, proteins) and processes (growth, heredity, and biochemical reactions), its deficiency will cause a slow growth and reproduction in plants (Costa et al., 2002, Razaq et al., 2017).

## 4. Natural and anthropogenic phenomenon yield fixed nitrogen

For plants to use N, it must be “*fixed*” or converted into biologically compatible, nitrogenous compounds such as nitrates (NO3), nitrites (NO2), and ammonium salts (NH3). Other than electrochemical (non-biological) fixation and N-fixing organisms (Rhizobium, Bacillus, and Clostridium carrying out biological fixation), fixed N input to the environment is also an important byproduct of livestock production via organic waste (Bouwman et al., 2013). Long before Haber-Bosh fertilizers became ubiquitous, poultry manure and other animal waste products (e.g., bat guano) were used as a source of supplemental N.

The burning of fossil fuels by electrical power-generating plants and internal combustion engines also perturbs natural N cycling (Fowler et al., 2013). In the Midwestern US, NOx produced as a by-product of energy production influences continental-scale N gradients as a function of distance from stationary source NOx emission sites (Butler et al., 2001, Elliot et al. 2007, Chapman et al., 2013). NOx emissions are oxidized to nitric acid (HNO3) and represent a major sink for soil deposition of N (Fowler et al., 2013).

## 5. Importance of the nitrogen cycle and its coupling with carbon cycle

Overall, cycling from atmospheric N2 to fixed forms and back into the atmosphere is central to the geochemistry of the earth. The microbially-mediated N fixation in the N cycle, in turn, is coupled to the global carbon cycle since fixed N is an essential element and often limiting for biological carbon fixation in primary production (Gruber and Galloway, 2008, Vries et al., 2016). Thus, N has the potential to influence biological carbon sinks for greenhouse gas. Anthropogenic input to the N cycle is schematized in Figure 3.

**Figure 3:**
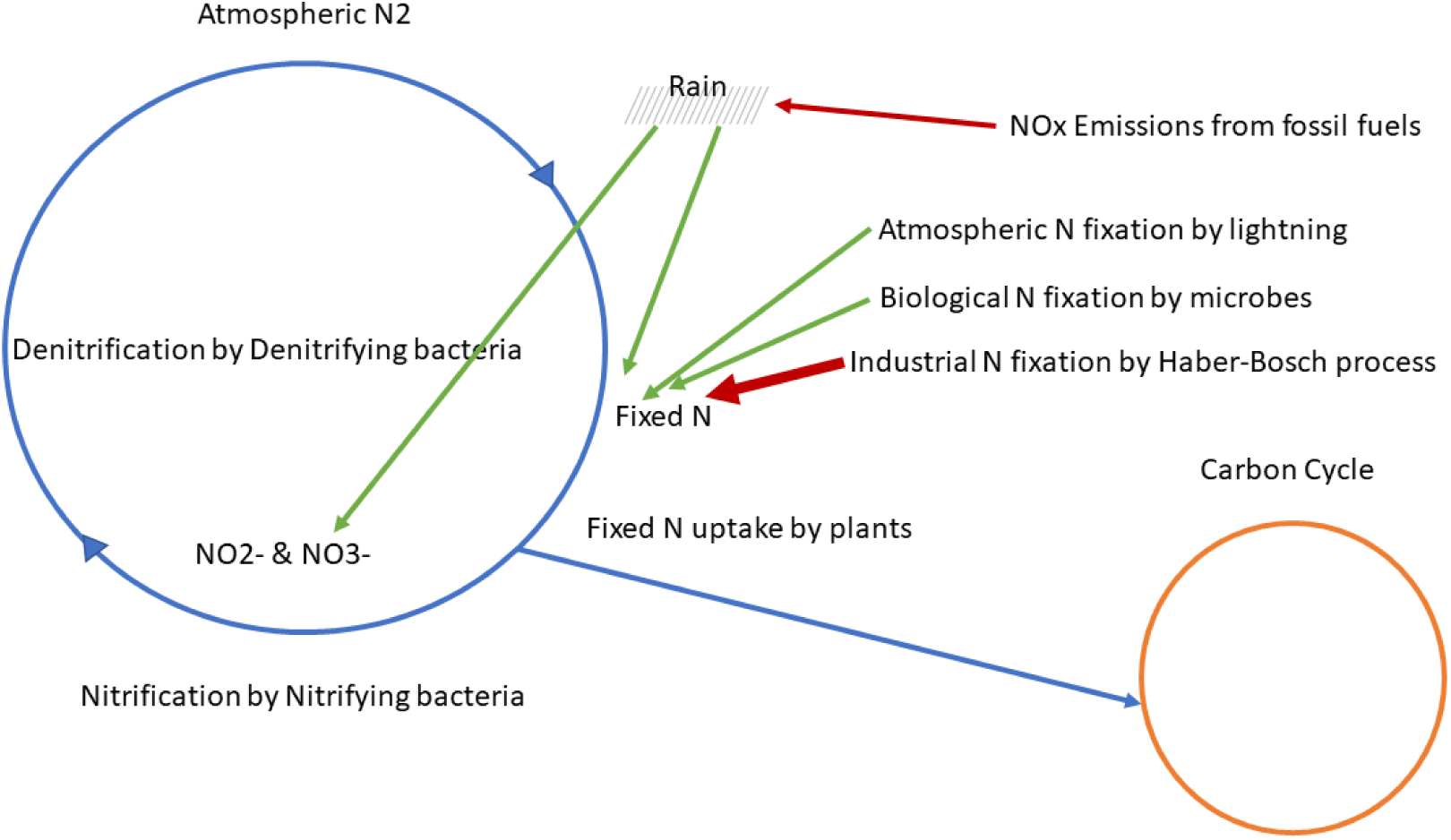
Anthropogenic inputs to the N Cycle. As can be seen, N flows through the biosphere and through abiotic systems in a loop that is coupled to the carbon cycle. Non-biological fixation occurs primarily via Haber-Bosh industrial production of fixed N (bold red), but also as a result of human energy production and consumption (thin red).

**Figure 4:**
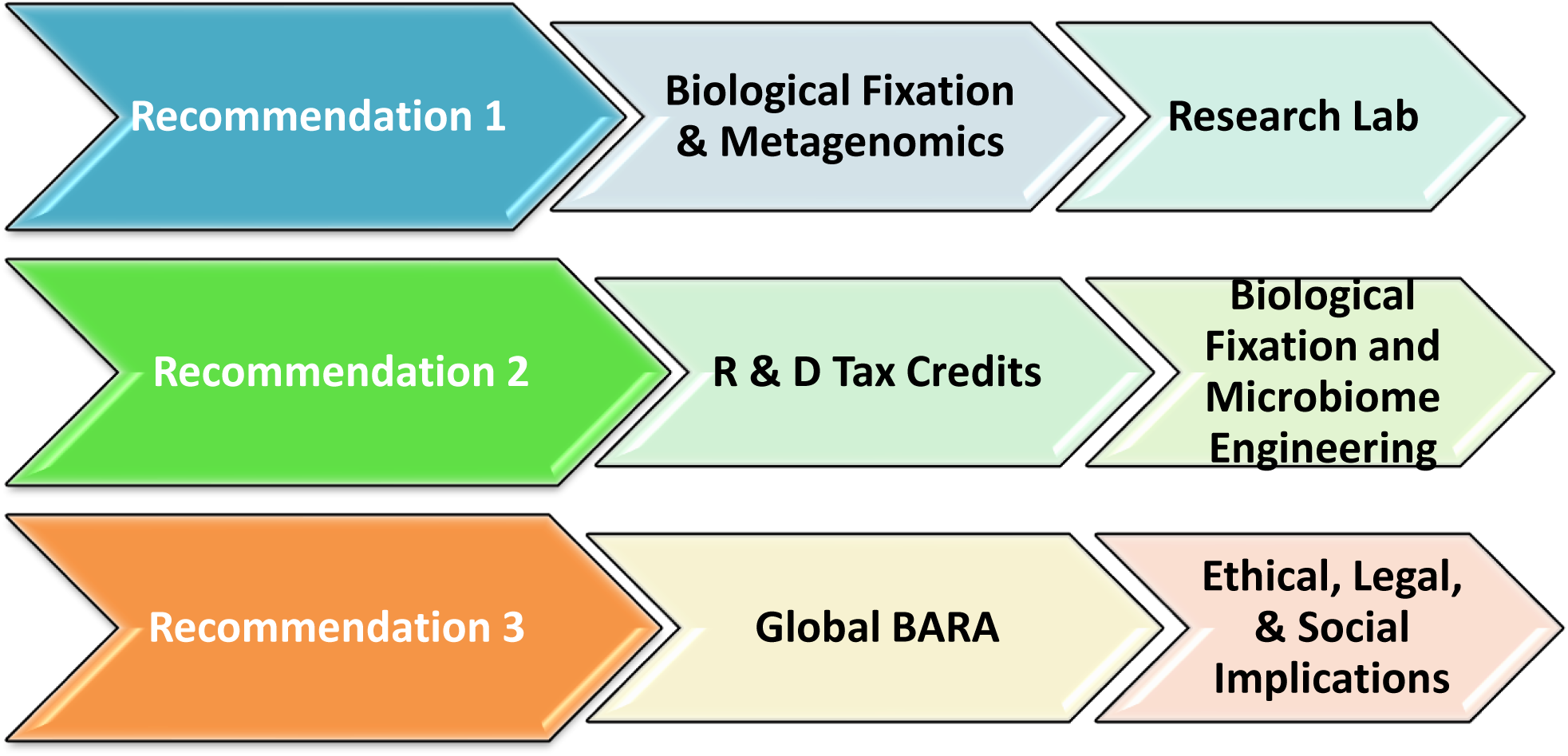
A coherent strategy for ensuring that future agriculture includes biological fixation as a prime strategy. The three recommendations presented in this perspective work together to ensure support for innovation and research into a Haber-Bosh free future agriculture.

## 6. Implications for the environment, microbes, and humans

Excess N supplementation to agricultural food systems along with other anthropogenic perturbations in the N cycle (figure 3) affect the biosphere and represent interesting avenues of study for life scientists. It has been shown that an inverse relationship exists between deposited N and plant diversity, determined from a long-term sampling of sites across the United States (Simkin et al., PNAS 2016). N has also been implicated in changes in coral reef microbiome diversity (Brock et al., 2018). Zhou et al. (2017) reported that experimentally added N to soy and wheat test plots in China caused profound changes in the soil microbiome diversity. Given the evidence for the coupling of microbiomes and host niche (Taş et al., 2018), it is vital to investigate how species composition will change as anthropogenic N input to the environment further increase over the coming decades.

In addition to negative impacts on biodiversity, it would be useful to investigate how antimicrobial resistance genes (ARGs) in microbiomes respond to increasing N supplementation. Subsequent findings will give an insight into selection pressure for ARGs and antibiotic-resistant microorganisms (antibiotic-resistant bacteria). This also has implications for human gut microbiome (HGM). The HGM is coupled to phytobiomes via food systems including crops and livestock, which are themselves tightly coupled. Thus, antibiotic resistance in food system microbiomes has the potential to produce the same in HGMs. Currently, antibiotic resistance represents a major health challenge across the globe, adding $20 billion in excess cost alone to the US health system (Naylor et al., 2018). As microbes increase their resistance to the current antibiotics, humans are increasingly at risk of developing infections for which there is no effective treatment. For instance, more than two million people per year acquire serious infections with bacteria resistant to one or more antibiotics in the US (CDC, 2013). Even in cases where treatment is available, the selection pressure of the treatment produces a trajectory across the microbial phenotypic landscape that only increases the problem.

## 7. What needs to be done?

Of many approaches, we advocate for microbiome-mediated N fixation, or simply *biological fixation* for a few reasons. The benefit of the approach is multifaceted. First, it is naturally available. Second, we believe it is replicable and scalable in many settings. Third, it preserves biodiversity and mitigates environmental hazards. The potential of biological fixation for agricultural cereal production is indicated by the recent finding of substantial N-fixing microbial associations in an indigenous landrace of maize (Van Deynze et al., 2018). For legumes, it is well known that microorganisms or microbiome (i.e., *Bradyrhizobium japonicum*) can persist in leguminous plants and assist them with N fixation (Kiers et al., 2003). The plants as a host, containing safe havens in their root nodules, provide microorganisms with a stable environment and nutrients needed to grow and persist. In turn, microorganisms offer the host with specific nutrients for growth and desirable traits such as antibiotic resistance for protection and other important mechanisms for reproduction (Hacquard et al., 2015; Philippot et al., 2013).

For biological fixation to happen in non-legume crops, there is a need for research that engineers novel microbiomes that can be grown and applied to food systems at scale. As early as the 1970’s, it has been suggested to engineer for biological fixation in non-legumes. Hardy and Havelka (1975) wrote that “cereals that could provide their own fertilizer are beyond doubt the biggest prize of all in the gift of the new biology.” Contemporaneous with that report, the N-fixing genes (*nif*) of *Klebsiella pneumoniae* had been inserted into *Escherichia coli*, causing the nonfixing bacterium capable of growing in the absence of combined N (Dixon & Postgate, 1972). Since then, there has been some research and attempts on introducing N fixation in nonfixing plants and transferring *nif* genes into heterologous hosts (Yang et al., 2018; Good, 2018; Burén, 2018; Allen et al., 2107; Pérez-González, 2017; López-Torrejón, 2016); however, scientists still lack a comprehensive understanding of the composition of the microorganisms that carry out N fixation processes. To better understand N fixation processes and other biological functions that microorganisms provide plants or their hosts, it is first essential to determine why a microorganism or community evolved, why it persists, and what benefits it imparts to the host itself, other organisms, and the community. Identification of signature microbes and community composition that can support biological N fixation in staple crop food systems is of the highest importance. Sensitive, comprehensive, and state of the art whole genome, shotgun metagenomics provides an approach to highlight such signature microbes and community composition. Using metagenomics, a comparative investigation of microbial samples from both legumes and non-legumes would help understand the relative changes in the function, genetic composition, and abundance of microbes.

After the descriptive and applied metagenomic research findings, the subsequent step would be to find simple and robust genetic components (or genes) capable of fixing N in cellular environments (e.g. in non-legumes) different from their natural hosts. Indeed, the resulting microbiome would have to be grown cheaply and at scale to ensure the viability and replicability of this solution.

With the above in mind, we make the following recommendations:

1. Identification of signature microbes that are useful for N fixation is a priority. Modern high throughput metagenomics provides an approach to highlight such signature microbes and community composition. We recommend that private industry partner with governments to establish a pre-competitive research laboratory equipped with the latest technologies that conduct the cutting-edge metagenomic research.
2. A solution as new as microbiome engineering would need government’s support for its identification, testing, and implementation. While there are many activities (including experimentation with new pesticides to control disease, new product development through cross-breeding, and design of new irrigation systems among other activities) that qualify for R & D tax credits, we recommend the U.S. Department of Agriculture (USDA) and state governments to provide R & D tax credits to ag-bio companies specifically geared towards the viability of biological fixation and microbiome engineering.
3. To coordinate investments and channelize knowledge gathered from the basic plant and agricultural sciences to agriculture in the field, we advocate the launch of a global agriculture innovation initiative, called Biological Advanced Research in Agriculture (BARA). Backed by the UN FAO, BARA would be an international scientific collaborative effort that would advance research on biological fixation among other aspects of agricultural innovation. The initiative would also address all the ethical, legal, and social implications around microbiome engineering. Needless to mention that without strong support of technologically advanced and some emerging economies such as the US, Russia, France, Germany, China, and India the initiative would be toothless.

## 8. Conclusion

With the increasing population and deteriorating environment, the excessive use of fertilizers cannot be ignored. Biological N fixation must be investigated and extended to the three staple crops (rice, wheat, and maize) that feed the world population. Apart from the environmental hazards of increasing fertilizer use, increased antibiotic resistance in human gut microbiome further makes the research about biological N fixation central. While current solutions are limited, we believe that a more fruitful approach for future agriculture in overcoming N challenge is to leverage biological N fixation as a practical and efficient strategy. The N challenge is a call for science to lead and innovate. A disruptive solution, such as microbiome engineering, can only be accomplished when scientists, industries, and governments consider it a high priority.

## Author contribution statement

SK, NK, and JO wrote the manuscript.

